# Dynamic O-GlcNAcylation of Sec23-interacting protein regulates COPII function

**DOI:** 10.64898/2025.12.18.695207

**Authors:** Tetsuya Hirata, Quyen Nguyen, Coco Liu, Michael Boyce

**Affiliations:** Department of Biochemistry, Duke University School of Medicine

**Keywords:** COPII, ER exit site, glycosylation, Sec23IP, Sec31A, O-GlcNAcylation, protein transport

## Abstract

About one-third of the eukaryotic proteome transits the secretory pathway to reach its correct cellular or extracellular destination. At the earliest stage, transport from the endoplasmic reticulum (ER) to the ER-Golgi intermediate compartment (ERGIC) or Golgi apparatus is mediated by coat protein complex II (COPII). COPII coats consist of inner and outer layers formed by Sec23–Sec24 heterodimers and Sec13–Sec31 heterotetramers, respectively, which initially assemble at ER exit sites (ERES) to form transport carriers. Sec23-interacting protein (Sec23IP) links the inner and outer coats through its interactions with both Sec23A and Sec31A, positioning it as a key potential regulator of COPII function. However, the mechanisms controlling Sec23IP activity remain poorly understood. Here, we investigate how physiological stimuli regulate COPII function through the dynamic modification of Sec23IP by O-linked β-*N*-acetylglucosamine (O-GlcNAc), a reversible, intracellular form of glycosylation. We first validated Sec23IP as a bona fide O-GlcNAcylated protein. Rescue experiments in Sec23IP knockout cells with a nearly unglycosylatable mutant protein demonstrated the essential role of O-GlcNAcylation in the intrinsically disordered domain in protein transport and in recruiting Sec31A to ERES. Moreover, O-GlcNAcylation of Sec23IP increased during protein transport, coinciding with a reduction in its interaction with Sec31A. These results indicate that distinct site-specific O-GlcNAcylation of Sec23IP spatiotemporally modulates its association with Sec31A to fine-tune ERES recruitment and COPII assembly/disassembly. Our work provides new insight into Sec23IP regulation and suggests that O-GlcNAc on other COPII proteins may govern carrier formation, uncoating, and transport.

## Introduction

In eukaryotic cells, roughly one-third of newly synthesized proteins enter the secretory pathway for targeting to specific intracellular locations or the extracellular space (1, 2). In humans, disruption of protein trafficking can lead to severe disorders, including neurological, skeletal, and hematological diseases, underlining its physiological importance (3–9). The endoplasmic reticulum (ER) is the entry point of the secretory pathway, where nascent polypeptides fold and mature. Properly folded secreted and transmembrane proteins are then concentrated at specialized domains known as ER exit sites (ERES), where transport towards the Golgi is initiated (1–4).

Coat protein complex II (COPII) plays a central role in ER-to-Golgi trafficking by recruiting cargo proteins and generating transport carriers (1–4, 10). The COPII coat comprises two subcomplexes: an inner coat of Sec23–Sec24 heterodimers and an outer coat of Sec13–Sec31 heterotetramers (1–4, 10). Once assembled, COPII carriers are transported to the ER–Golgi intermediate compartment (ERGIC) or the Golgi. Recent imaging studies have revealed that, in addition to discrete vesicles (11), COPII transport may occur via tubules or tunnels emanating from ERES, with COPII proteins localized at the neck regions (12, 13), though this model is controversial (14). While the biochemical and structural foundations of COPII-mediated transport have been extensively characterized (12, 13, 15–23), the dynamic regulation of COPII function in response to cellular or physiological cues remains largely enigmatic.

The essential components of COPII coats were established through *in vitro* reconstitution experiments, showing that Sar1, Sec23, Sec24, Sec13, and Sec31 are sufficient to generate COPII vesicles (15). Additional accessory proteins, including Sec16 and Sec23-interacting protein (Sec23IP, also known as p125A), have since been identified as critical regulators of ER export in live mammalian cells (19, 24). Sec23IP was originally characterized as a 125-kDa Sec23A-binding protein containing a phospholipase A1 homology domain, though enzymatic activity has not been detected (25, 26). Subsequent studies revealed that Sec23IP also interacts with Sec31A and is essential for conventional COPII-mediated transport in mammalian cells (24). More recently, Sec23IP has been implicated in specialized COPII-dependent trafficking processes, such as those that transport large cargoes like collagens via interactions between Sec23IP and the ERGIC-localized vacuolar protein sorting 13 homolog B and phosphatidylinositol 4-phosphate (27–29).

The molecular mechanisms governing Sec23IP activity and its interactions with COPII components remain incompletely understood, but post-translational modifications likely play a role. For example, high-throughput mass spectrometry (MS) analyses indicated that Sec23IP is heavily glycosylated with O-GlcNAc (O-linked β-*N*-acetylglucosamine), a reversible, highly conserved monosaccharide modification of intracellular proteins (30–37). Over 5,000 human proteins are reportedly modified with O-GlcNAc at 7,000 distinct sites (38). O-GlcNAcylation is carried out by the nuclear-cytoplasmic glycosyltransferase OGT, transferring GlcNAc moieties onto Ser/Thr residues, and is removed by the glycoside hydrolase OGA (35, 36, 39). This rapid reversibility allows O-GlcNAc to regulate various cellular processes, including COPII trafficking (39–43). We have previously identified O-GlcNAcylation sites on multiple COPII components, including Sec23A, Sec31A, Sec24C, and Sec24D (44, 45). Subsequently, we and others investigated the function of this modification on Sec23A, Sec24D, and Sec31A (44, 46), revealing that site-specific O-GlcNAcylation regulates protein-protein interactions required for collagen transport (45, 47). Moreover, we showed that glucose fluctuations, cell cycle progression, and collagen transport effect dynamic O-GlcNAcylation changes on Sec31A, Sec24C, and Sec24D, suggesting that O-GlcNAc is a central regulator of COPII function in response to diverse stimuli (44, 47, 48). Here, we extend these studies by addressing how Sec23IP O-GlcNAcylation impacts COPII function.

## Results

### Validation of Sec23IP O-GlcNAcylation

Previous MS studies indicated that Sec23IP is O-GlcNAcylated (30–34), but these results had not been independently confirmed. To validate prior MS data, we expressed Myc-6xHis-Sec23IP in HEK293 cells, subjected them to pharmacological OGT or OGA inhibition via the small molecule Ac_4_5SGlcNAc (5SG) (49) or Thiamet-G (TG) (50), respectively, and analyzed the protein via immunoprecipitation and Western blot (IP/WB). O-GlcNAcylation was assessed with three monoclonal antibodies – 18B10, RL2, and 9D1 – which recognize partially overlapping subsets of O-GlcNAcylated epitopes (51). All three antibodies detected O-GlcNAcylated Sec23IP in vehicle-treated controls, with the signal decreasing upon 5SG treatment and strongly increasing upon TG treatment (Fig. 1A). RL2 and 9D1 also detected O-GlcNAcylation on endogenous Sec23IP in HeLa and HEK293 lysates (Fig. 1B). Next, we generated Sec23IP knockout (KO) HeLa and HEK293 cells using CRISPR–Cas9 (Fig. S1A, left and middle) and performed IPs with anti-Sec23IP. No signal was detected in KO cells (Fig. 1B), confirming antibody specificity. Collectively, these data demonstrate that Sec23IP is a bona fide and dynamically O-GlcNAcylated protein.

**Figure 1.**
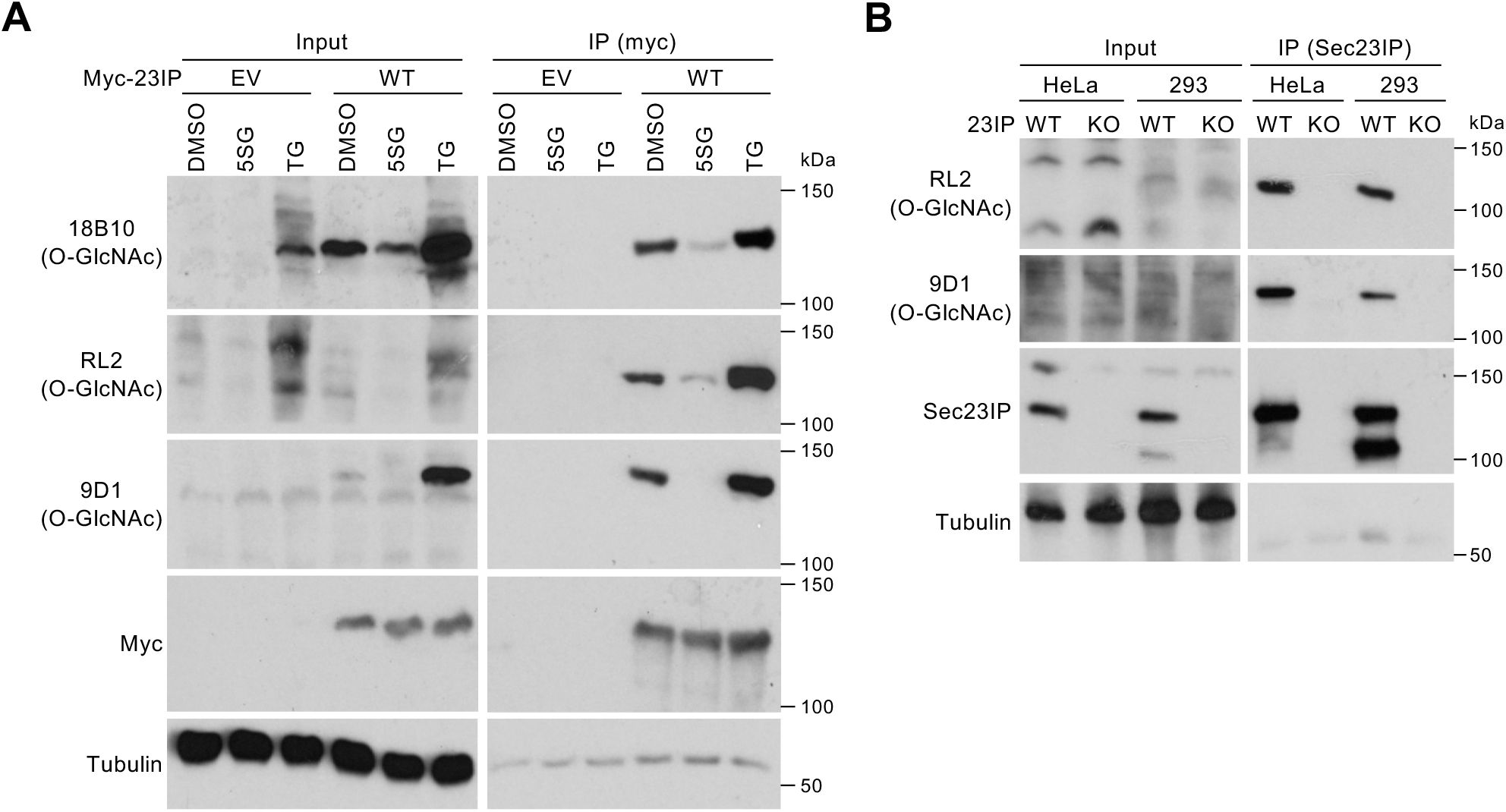
Human Sec23IP is extensively O-GlcNAcylated. A. Myc-6xHis-tagged human Sec23IP (Myc-23IP) was expressed in HEK293 cells. Cells were treated with DMSO vehicle, 50 µM Ac_4_5SGlcNAc (5SG, OGT inhibitor), or 50 µM Thiamet-G (TG, OGA inhibitor) for 24 hr. Myc-23IP was IP-ed with anti-myc antibody and analyzed by WB. n = 3, representative images shown. B. Endogenous Sec23IP was IP-ed with anti-Sec23IP antibody from WT or Sec23IP (23IP)-KO HeLa and HEK293 (293) cells and analyzed by WB. n = 2, representative images shown.

### O-GlcNAcylation of the Sec23IP IDR Is Critical for Efficient ER-Golgi Transport

Sec23IP has been implicated in conventional and unconventional transport of both small cargoes (e.g., vesicular stomatitis virus glycoprotein, VSVG) (24) and bulky cargoes such as collagen (28, 29). Therefore, we evaluated its contribution to these pathways in functional experiments. Conventional transport was assessed using E-Cadherin as a well-established model cargo (47, 52, 53) in a retention using selective hooks (RUSH) assay (54). Consistent with prior studies (47, 53), in wild type (WT) cells, E-Cadherin accumulated in the ER at time zero and efficiently moved to the Golgi over time after release of the RUSH probe by biotin addition (Fig. S1B-C). In contrast, Sec23IP-KO HeLa cells displayed only partial Golgi localization of the E-Cadherin RUSH probe even after 40 min, indicating impaired transport (Fig. S1B-C).

To test bulky cargo transport, we used human chondrosarcoma SW1353 cells, which express endogenous collagen and rely on specialized large COPII carriers (45, 47, 55). We generated Sec23IP-KO SW1353 cells (Fig. S1A, right) and used a collagen transport assay involving a 40 °C block followed by a shift to permissive 32 °C with ascorbate addition, which promote collagen maturation and ER exit (56). Unlike E-Cadherin, collagen transport was comparable between WT and KO cells (Fig. S1D-E), suggesting that Sec23IP is not essential for collagen trafficking in this context.

To determine how O-GlcNAcylation contributes to Sec23IP function, we queried the O-GlcNAc database (57) and identified 22 reported glycosites. Interestingly, 21 of these lie within the N-terminal intrinsically disordered region (IDR), whose function remains mysterious (Fig. 2A and Fig. S2). We generated an O-GlcNAc-deficient mutant in which all N-terminal IDR glycosites were mutated to alanine (“NA” mutant). NA expressed at levels comparable to WT (Fig. 2B) and localized normally to Sec16A-marked ERES (Fig. 2C), indicating proper folding. However, the number of Sec23IP and Sec16A puncta increased significantly in NA-expressing cells, relative to WT (Fig. 2C-D), suggesting a transport defect leading to ERES accumulation.

**Figure 2.**
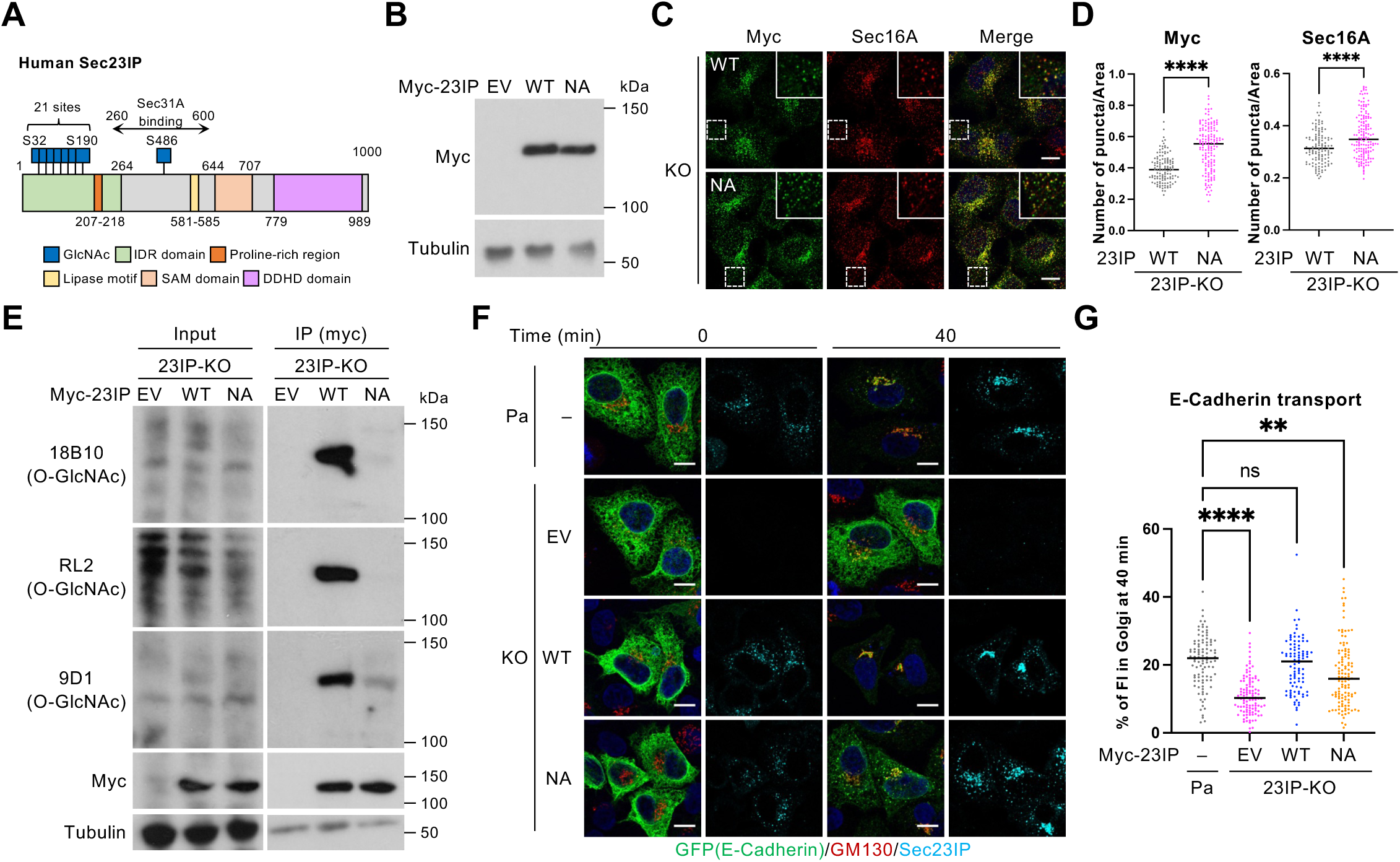
O-GlcNAcylation of the Sec23IP IDR domain is required for conventional COPII-dependent protein transport. A. Domain structure and potential O-GlcNAc sites of human Sec23IP. Previous high-throughput MS analyses identified 22 potential O-GlcNAc sites on human Sec23IP, 21 of which are located on its N-terminal IDR. Reported O-GlcNAc sites on human Sec23IP are depicted in Fig. S2. B. HEK293 cells were transfected with empty vector (EV) or expression constructs encoding Sec23IP WT or NA mutant for 24 hr and analyzed by WB. n = 2, representative images shown. C. HeLa-23IP-KO cells were transfected with expression constructs encoding Sec23IP WT or NA mutant for 24 hr and analyzed by IF. n = 3, representative images shown. Sec16A: ERES marker. Scale bars: 10 µm. D. Quantification of experiments depicted in C. Left: Number of Myc-23IP puncta per cell area (n = 3). Right: Number of Sec16A puncta per cell area (n = 3). Data were analyzed by Welch’s *t*-test. **** *p* < 0.0001. E. HEK293-23IP-KO cells were transfected with expression constructs encoding Sec23IP WT or NA mutant for 24 hr. Myc-23IP was IP-ed with anti-myc antibody and analyzed by WB. n = 3, representative images shown. F. Parental HeLa and 23IP-KO cells were co-transfected with plasmids encoding the E-cadherin RUSH system plus either nothing (–), empty vector (EV), or Sec23IP WT or NA mutant for 24 hr. For RUSH assays, cells were incubated with 50 µM biotin and 100 µg/ml cycloheximide for the indicated times and analyzed by IF for E-cadherin (GFP), GM130 (Golgi marker), and Sec23IP. n = 3, representative images shown. Scale bars: 10 µm. G. Quantification of experiments depicted in F. Percentage of GFP FI co-localizing with the Golgi (GM130) was calculated (n = 3). Data were analyzed by one-way ANOVA with *post hoc* Tukey test. ** *p* < 0.01; **** *p* < 0.0001; ns: not significant.

O-GlcNAcylation of NA was nearly abolished, compared to WT, as judged by IP/WB (Fig. 2E), confirming extensive glycosylation of the IDR in live cells. To examine the functional consequences of loss of Sec23IP O-GlcNAcylation, we re-expressed WT or NA Sec23IP in Sec23IP-KO cells and quantified E-Cadherin RUSH transport. As before, KO cells displayed severe transport delays (Fig. 2F-G). WT Sec23IP fully rescued transport, whereas NA only partially restored it, with significantly reduced Golgi-localized E-Cadherin (Fig. 2F-G). These data indicate that O-GlcNAcylation of the IDR is essential for Sec23IP function in the COPII-dependent transport of a canonical model cargo.

### O-GlcNAcylation of the Sec23IP IDR Regulates its Interaction with Sec31A

Because Sec23IP reportedly binds Sec23A and Sec31A (24, 25), we next investigated whether O-GlcNAcylation influences these interactions. WT or NA Sec23IP was expressed in Sec23IP-KO cells and subjected to co-IP. Interaction with Sec23A was similar for WT and NA, but the interaction with Sec31A was significantly reduced in the NA mutant (Fig. 3A-B). To test the impact on the subcellular localization of Sec31A, we performed immunofluorescence (IF) analysis. Sec23IP-KO cells displayed a marked reduction in the mean fluorescence intensity (MFI) of Sec31A puncta, relative to control cells (Fig. S3A-B), despite normal Sec31A expression levels overall (Fig. S3C-D). These observations are consistent with reduced recruitment of Sec31A to ERES in the absence of functional Sec23IP (24). Next, we complemented the KO cells with either WT or NA Sec23IP. Consistent with our co-IP results, WT Sec23IP restored Sec31A puncta in Sec23IP-KO cells but NA did not (Fig. 3C-D), confirming that IDR O-GlcNAcylation specifically supports Sec31A recruitment to ERES.

**Figure 3.**
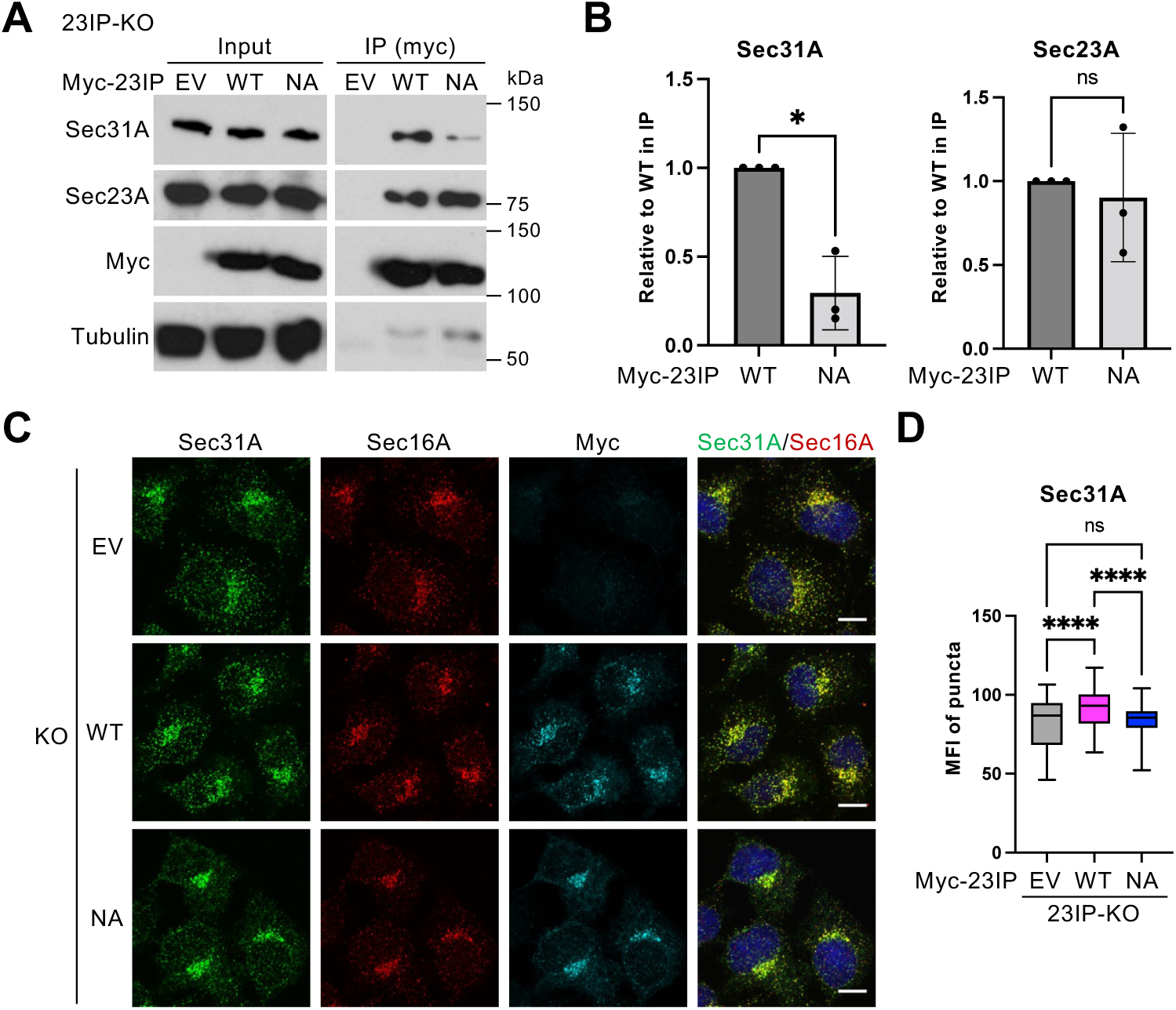
O-GlcNAcylation of the Sec23IP IDR domain is required for Sec31A recruitment to ERES. A. HEK293-23IP-KO cells were transfected with empty vector (EV) or expression constructs encoding Sec23IP WT or NA mutant for 24 hr. Myc-23IP was IP-ed with anti-myc antibody and analyzed by WB. n = 3, representative images shown. B. Quantification of experiments depicted in A. Signal from LI-COR Sec31A or Sec23A blots was normalized to myc signal, and intensities relative to WT were calculated (n = 3). Data were analyzed by Welch’s *t*-test. * *p* < 0.05; ns: not significant. C. HeLa-23IP-KO cells were transfected with empty vector (EV) or expression constructs encoding Sec23IP WT or NA mutant for 24 hr and analyzed by IF. n = 3, representative images shown. Sec16A: ERES marker. Scale bars: 10 µm. D. Quantification of experiments depicted in C. MFI of Sec31A puncta was calculated and plotted (n = 3). Data were analyzed by one-way ANOVA with *post hoc* Tukey test. **** *p* < 0.0001; ns: not significant.

### Sec23IP O-GlcNAcylation Is Dynamically Regulated During Protein Transport

COPII formation is regulated by cargo flux (1–4), and we previously showed that the cargo-recruiting subunit Sec24D becomes O-GlcNAcylated in response to collagen ER egress (47). We therefore asked whether Sec23IP O-GlcNAcylation is similarly regulated. Because E-Cadherin transport is affected in Sec23IP-KO HeLa cells (Fig. 2F-G and Fig. S1B-E), we examined the O-GlcNAc changes upon RUSH E-Cadherin trafficking in Sec23IP-KO cells expressing myc-6xHis-Sec23IP. O-GlcNAcylation of Sec23IP significantly increased over time during transport, as measured by the 9D1 antibody (Fig. 4A-B). A similar trend was observed in RL2 blots, though it was not statistically significant, whereas 18B10 signal remained essentially unchanged (Fig. 4A-B). These data demonstrate site-specific, dynamic regulation of Sec23IP O-GlcNAcylation during cargo trafficking. Co-IP revealed that the interaction of Sec23IP with Sec31A decreased markedly at 20 min (Fig. 4C-D), coinciding with increased O-GlcNAcylation. By contrast, the Sec23IP-Sec23A interaction was unaffected under the same conditions (Fig. 4C-D), consistent with our observation that the Sec23IP-Sec23A interaction was independent of Sec23IP glycosylation (Fig. 3A-B). Finally, IF showed that Sec31A distribution remained stable during transport, whereas Sec23IP puncta intensity increased and puncta number decreased over time (Fig. 4E-F). All together, these results show that protein transport induces spatiotemporally specific O-GlcNAcylation of Sec23IP, which inhibits its interaction with Sec31A.

**Figure 4.**
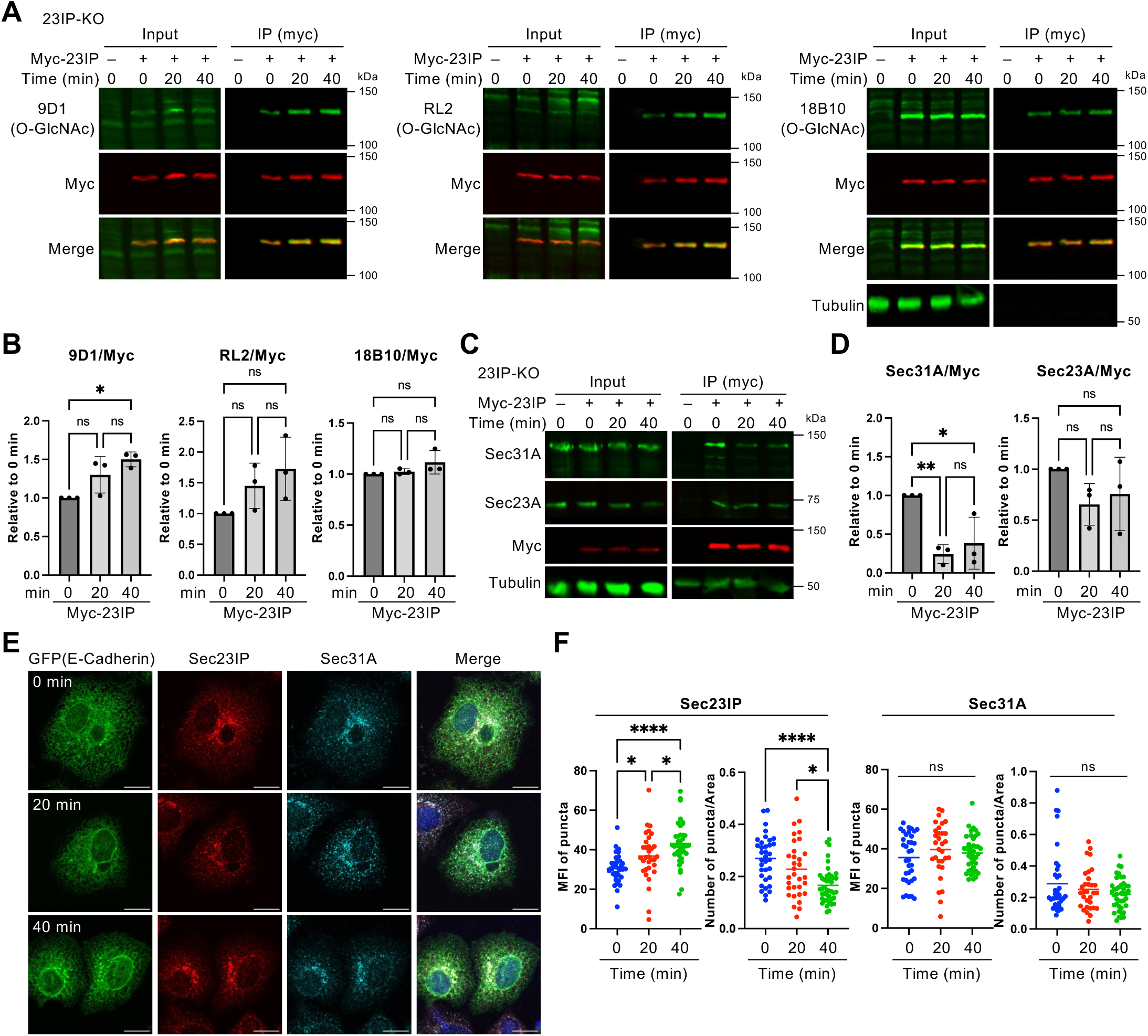
O-GlcNAcylation of Sec23IP is dynamically regulated in response to COPII protein transport. A. HEK293-23IP-KO cells were transfected with empty vector (EV) or expression constructs encoding Sec23IP WT for 24 hr, and myc-23IP was IP-ed with anti-myc antibody at 0, 20, or 40 minutes after induction of E-Cadherin transport and analyzed by WB. n = 3, representative images shown. B. Quantification of experiments depicted in A. Signal from O-GlcNAc blots of purified Sec23IP was normalized to myc signal, and intensities relative to time-zero were calculated (n = 3). Data were analyzed by one-way ANOVA with *post hoc* Tukey test. * *p* < 0.05; ns: not significant. C. HEK293-23IP-KO cells were transfected with empty vector (EV) or expression constructs encoding Sec23IP WT for 24 hr, and myc-23IP was IP-ed with anti-myc antibody at 0, 20, or 40 minutes after induction of E-Cadherin transport and analyzed by WB. n = 3, representative images shown. D. Quantification of experiments depicted in C. Signal from Sec31A or Sec23A blots was normalized to myc signal, and intensities relative to time-zero were calculated (n = 3). Data were analyzed by one-way ANOVA with *post hoc* Tukey test. * *p* < 0.05; ** *p* < 0.01; ns: not significant. E. Parental HeLa cells expressing the E-cadherin RUSH system were incubated with 50 µM biotin and 100 µg/ml cycloheximide for the indicated times and analyzed by IF for E-cadherin (GFP), Sec23IP, and Sec31A. n = 3, representative images shown. Scale bars: 15 µm. F. Quantification of experiments depicted in E. MFI of Sec23IP or Sec31A puncta and the number of Sec23IP or Sec31A puncta per cell area were calculated and plotted (n = 3). Data were analyzed by one-way ANOVA with *post hoc* Tukey test. * *p* < 0.05; **** *p* < 0.0001; ns: not significant.

## Discussion

Because Sec23IP interacts with both the inner and outer COPII coats, it is well-positioned to influence trafficking. However, its upstream regulation and downstream functions have remained largely unclear. Here, we demonstrate that dynamic O-GlcNAcylation of Sec23IP is essential for recruiting Sec31A (but not Sec23A) to ERES, thereby facilitating COPII carrier biogenesis. Furthermore, Sec23IP O-GlcNAcylation increases over the course of cargo transport, correlating with a loss of Sec31A interaction. Together, these data suggest a model in which glycosylation at some constitutive Sec23IP site(s) potentiates initial outer coat recruitment to ERES, whereas later, inducible glycosylation events at distinct sites promote COPII carrier uncoating (Fig. 5).

**Figure 5.**
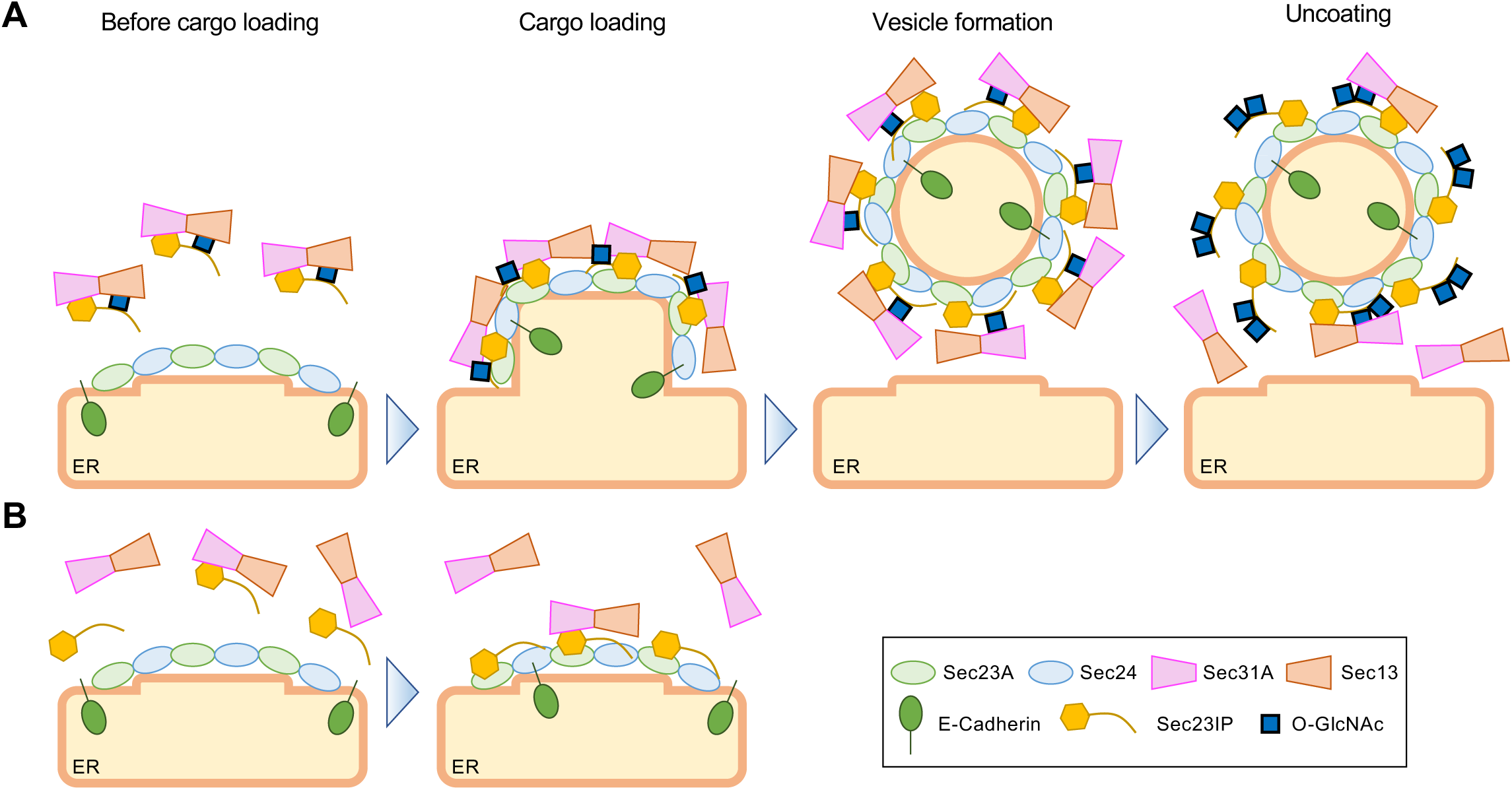
Proposed model for functional regulation of COPII trafficking by spatiotemporally dynamic Sec23IP O-GlcNAcylation. A. At steady state, the Sec23IP IDR domain is constitutively O-GlcNAcylated on specific sites, which promotes the formation of a ternary complex with the Sec31A-Sec13 outer coat of COPII. When protein transport is initiated (i.e., by endogenous protein transport under physiological condition or biotin addition in RUSH experiments), conventional cargo proteins (e.g., E-Cadherin) are loaded into ERES via direct or indirect interaction with Sec24, which in turn recruits a ternary Sec23IP-Sec31A-Sec13 complex, generating COPII carriers. After transport is underway, the Sec23IP IDR is further O-GlcNAcylated at distinct, inducible sites, promoting carrier uncoating. In each case, the specific O-GlcNAc site(s) involved remain to be identified. B. In the Sec23IP NA mutant, constitutive O-GlcNAcylation is greatly diminished and ternary complex formation with Sec31A-Sec13 is inhibited, reducing outer coat recruitment to ERES or pre-budding complexes in response to cargo demand and delaying transport.

Although Sec23IP had been reported as O-GlcNAcylated in MS screens previously (30–34), no study had validated these results via orthogonal methods. We provide confirmatory evidence through IP/WB of expressed and endogenous protein, demonstrating recognition of Sec23IP by multiple O-GlcNAc antibodies, with signals altered by OGT/OGA inhibitors as expected (Fig. 1). Furthermore, removing all reported IDR O-GlcNAc sites in the NA mutant nearly abolished the modification (Fig. 2E). Residual anti-O-GlcNAc immunoreactivity of NA is likely due to additional sites, such as S486 (30). Comprehensive identification of all physiological Sec23IP O-GlcNAc sites will require systematic MS analysis of purified endogenous protein in future studies.

Using the NA mutant, we showed that IDR O-GlcNAcylation is required for Sec23IP’s interaction with Sec31A but not Sec23A (Fig. 3A-B). Consistent with this result, NA only partially rescued transport in Sec23IP-KO cells (Fig. 2F-G), likely due at least in part to impaired Sec31A recruitment (Fig. 3C-D). Given these observations, coupled with prior reports that Sec23IP forms a ternary complex with Sec31A and Sec13 in the cytosol (24), we propose that constitutive O-GlcNAcylation of specific sites in the Sec23IP IDR positively regulates the early stage of COPII transport by recruiting outer coat components to ERES and pre-budding complexes to facilitate carrier biogenesis (Fig. 5).

By combining RUSH assays and IP, we demonstrated that Sec23IP O-GlcNAcylation increases during E-Cadherin trafficking (Fig. 4A-B), most likely within its IDR, where nearly all reported sites reside (Fig. 2A, 2E). Although we previously observed dynamic O-GlcNAcylation on Sec24D during collagen transport (47), the current report is, to our knowledge, the first demonstration of temporal O-GlcNAc changes during transport of a smaller, more typical COPII cargo. Together, these data suggest that dynamic O-GlcNAcylation cycling may be a key modulator of COPII-dependent protein trafficking for a range of cargoes. The Sec23IP-Sec31A interaction diminished as Sec23IP O-GlcNAcylation increased (Fig. 4C-D), and Sec23IP puncta MFI rose in concert (Fig. 4E-F), suggesting that Sec23IP glycosylation at this stage negatively regulates Sec31A association. Therefore, we propose that distinct Sec23IP glycosites are inducibly modified later during transport, inhibiting Sec31A binding and promoting carrier uncoating (Fig. 5). As Sec31A puncta number and intensity did not change under these conditions (Fig. 4E-F), we hypothesize that the released outer coat components may be rapidly recycled and re-recruited to ERES for further rounds of cargo exports. In future studies, it will be important to test this model by identifying the specific Sec23IP residues where O-GlcNAcylation increases during transport and testing the effects of unglycosylatable mutations at those sites in functional trafficking and Sec31A localization assays.

Notably, four reported O-GlcNAc sites in the IDR of human Sec23IP (T41, S111, T123, and S190) are conserved across vertebrates, and four others (S32, S87, S90, and T186) are conserved among mammals and either fish or amphibians (Fig. S2). COPII transport is fundamental to all eukaryotes, and our prior results indicate that the O-GlcNAc-dependent regulation of COPII is likely conserved at least from humans to fish (45, 47). Therefore, it is interesting to speculate that these conserved O-GlcNAc sites in Sec23IP could be functionally significant as constitutive and/or inducible glycosites. Further mutagenesis and trafficking analyses will be important future goals to test this hypothesis.

The biophysical basis for the O-GlcNAc-mediated regulation of the Sec23IP-Sec31A interaction remains as-yet unknown. The Sec23IP IDR O-GlcNAc sites are outside of the region reported to interact with Sec31A (Fig. 2A) (24). Therefore, it is unlikely that Sec31A directly binds to O-GlcNAc moieties on Sec23IP. Instead, constitutive IDR O-GlcNAcylation may influence the overall structure of Sec23IP, increasing its affinity toward Sec31A. Alternatively, because IDRs often participate in weak, multivalent interactions and liquid-liquid phase separation (LLPS) (58–61) and because COPII-related LLPS has been reported (2, 62–64), O-GlcNAcylation may modulate an LLPS-mediated recruitment mechanism. Indeed, O-GlcNAcylation has been reported to modulate LLPS formation in other contexts (65–68). Intriguingly, we recently discovered a similar control of protein-protein interactions by O-GlcNAcylation of the IDR of Sec24D as well (47), implying that this may be a general mode of COPII regulation. Structural studies and LLPS assays using O-GlcNAcylated IDR constructs will be required to test this notion.

E-Cadherin transport was strongly impaired in Sec23IP-KO HeLa cells, consistent with the known requirement for Sec23IP in the conventional transport of other model cargoes, such as VSVG (24). However, unlike recent reports implicating Sec23IP in collagen transport (28, 29), we did not observe similar defects in Sec23IP-KO SW1353 cells (Fig. S1D-E). Differences in species, cell type, or gene depletion strategies may account for this ostensible discrepancy. For example, KO cells may undergo compensatory adaptation, such as altered gene expression, which could obscure functional roles for Sec23IP in collagen trafficking.

In summary, we have identified dynamic O-GlcNAcylation of the Sec23IP IDR as an important regulator of COPII function. Our findings provide new insight into how COPII coat assembly is modulated and highlight Sec23IP O-GlcNAcylation as a key potential regulatory mechanism. We expect that future dissection of individual glycosites on Sec23IP and other COPII proteins will deepen our understanding of protein transport in health and disease.

### Experimental procedures

#### Antibodies and reagents

The following antibodies were used: Rabbit anti-collagen I antiserum (LF-68) was a gift from L. Fisher, National Institute of Dental and Craniofacial Research; Rabbit polyclonal anti-Sec23A (#8162) from Cell Signaling Technology; Mouse anti-c-Myc (9E10, #626802) and mouse anti-O-GlcNAc (RL2, #677902) from BioLegend; Mouse anti-O-GlcNAc (18B10.C7, #MA1-038), and mouse anti-O-GlcNAc (9D1.E4, #MA1-039) from ThermoFisher Scientific; Mouse anti-α-Tubulin (#T6074), and rabbit polyclonal anti-Sec23IP (#HPA038403) from Sigma-Aldrich; Rabbit polyclonal anti-Sec16A (#A300-648A) from Bethyl Laboratories; Mouse anti-Sec31A (32, #612351) and mouse anti-GM130 (35, #610822) from BD Biosciences; Chicken anti-GFP (#ab13970), and rat anti-myc (9E10, #ab206486) from Abcam; Rabbit anti-Sec23IP from Abcepta (#AP14511b); Goat horseradish peroxidase (HRP)-conjugated anti-mouse IgG (#1030-05), and goat HRP-conjugated anti-rabbit IgG (#4030-05) from SouthernBiotech; Goat IRDye 800CW-conjugated anti-mouse IgG (#925-32210), and goat IRDye 800CW-conjugated anti-rabbit IgG (#925-32211) from LI-COR; Goat Alexa Fluor 488-conjugated anti-mouse IgG (A-11001), goat Alexa Fluor 488-conjugated anti-rabbit IgG (A-11008), goat Alexa Fluor 488-conjugated anti-chicken IgY (A-11039), goat Alexa Fluor 594-conjugated anti-mouse IgG (A-11005), goat Alexa Fluor 594-conjugated anti-rabbit IgG (A-11012), goat Alexa Fluor 647-conjugated anti-mouse IgG (A-21235), goat Alexa Fluor 647-conjugated anti-rabbit IgG (A-32733), and goat Alexa Fluor 647-conjugated anti-rat IgG (A-21247) from ThermoFisher Scientific.

The following reagents and kits were used: Lipofectamine 3000 (Cat#L3000015) from ThermoFisher Scientific; Biotin (#B4501-100MG) from Sigma-Aldrich; Thiamet-G (TG) from Cayman Chemical (#13237).

Peracetylated 5S-GlcNAc (5SG) was synthesized by the Duke Small Molecule Synthesis Facility as described previously (49).

#### Plasmid construction

A single guide RNA (sgRNA) for human *SEC23IP* knock-out (5’-AGGCCCAGGGTTGTAAAGCG-3’) was subcloned into pSpCas9(BB)-2A-GFP (pX458, Addgene #48138) by the Duke Functional Genomics Facility. To obtain full length cDNA of human Sec23IP, pCMV-SPORT6-hSec23IP was purchased from DNAFORM (Clone ID: 6514739). To construct pLenti6-myc-6xHis-hSec23IP, full length cDNA of hSec23IP was amplified with a primer set hSec23IP-F: 5’-CTCGGATCCACTAGTgccgagagaaaacctaacggtg-3’ and hSec23IP-R: 5’-gaaagctgggtCGGCtcaatgctggggctgttctg-3’, and a pLenti6 backbone was amplified with primer sets pLenti-Gib-F2: 5’-cattggtaactgtcagaccaagtttactc-3’ and pLenti-hSec23IP-R: 5’-aggttttctctcggcACTAGTGGATCCGAGCTCGGTAC-3’, and pLenti-Sec23IP-F: 5’-cagccccagcattgaGCCGacccagctttcttgtac-3’ and pLenti-Gib-R2: 5’-gagtaaacttggtctgacagttaccaatg-3’. Amplified sequences were assembled with NEBuilder HiFi DNA Assembly Master Mix (New England BioLabs, #M5520AA) by mixing them at 2:1:1 ratio at 50 °C for 1 h. To construct plasmids expressing O-GlcNAc-deficient human Sec23IP mutants (NA), a gene block encoding the N-terminus of hSec23IP with mutations in the potential 21 O-GlcNAc sites were designed (5’-ATGGCCGAGAGAAAACCTAACGGTGGCAGCGGCGGCGCCTCCACTTCCTCATCGGGCACTAACTTACTTTTCTCCTCCTCGGCCACGGAGTTCgcCTTCAATGTGCCCTTCATCCCAGTCgCCCAGGCCgCCGCTTCTCCGGCCgCCCTGCTCTTACCGGGAGAGGATTCCACAGATGTTGGTGAGGAGGACAGCTTCCTTGGTCAGACTTCTATTCACACATCTGCCCCACAGACATTTAGTTACTTCTCTCAGGTAgCAgcCAGCgcTGATCCTTTTGGGAATATTGGACAGTCACCATTAACAgCTGCAGCAgCCgCAGTTGGACAAgCAGGATTCCCCAAGCCCCTGgCTGCTCTCCCTTTTgCAgCTGGAgCCCAAGATGTCgCGAATGCATTTgCACCAgCCATTgCGAAGGCTCAACCTGGTGCTCCACCTTCCTCACTGATGGGAATAAATTCTTATCTGCCTTCTCAGCCAAGTAGTCTCCCTCCTTCATATTTTGGGAACCAACCCCAAGGAATTCCCCAACCAGGATACAATCCATATCGCCATgCCCCTGGCAGCgcCAGGGCTAATCCTT AC-3’) and was amplified with a primer set hSec23IP-F: 5’-CTCGGATCCACTAGTgccgagagaaaacctaacggtg-3’ and hSec23IP-NA-R: 5’-GTAAGGATTAGCCCTGgcGCTGCCAGGGGcATG-3’. A pLenti6 backbone was amplified with primer sets pLenti-Gib-F2: 5’-cattggtaactgtcagaccaagtttactc-3’ and pLenti-hSec23IP-R: 5’-aggttttctctcggcACTAGTGGATCCGAGCTCGGTAC-3’, and hSec23IP-R191-F: 5’-gCCCCTGGCAGCgcCAGGGCTAATCCTTACATTGCACC-3’ and pLenti-Gib-R2: 5’-gagtaaacttggtctgacagttaccaatg-3’. Amplified sequences were assembled with NEBuilder HiFi DNA Assembly Master Mix by mixing them at 2:1:1 ratio at 50 °C for 1 h.

#### Cell culture

HEK293, HeLa and SW1353 cells were obtained from ATCC. HEK293, HEK293-Sec23IP-KO, HeLa, HeLa-Sec23IP-KO, SW1353, and SW1353-Sec23IP-KO cell lines were cultured in Dulbecco’s modified Eagle’s medium (DMEM) supplemented with 10% fetal bovine serum and 1% penicillin/streptomycin at 37 °C under 5% v/v CO_2_ conditions.

#### Inhibitor treatment

Cells were treated with 50 µM 5SG (OGT inhibitor) or 50 µM TG (OGA inhibitor) for 24 h at 37 °C.

#### Plasmid transfection

Cells at approximately 50% confluence grown on 6-cm or 15-cm dishes were transfected with each plasmid using Lipofectamine 3000 in accordance with the manufacturer’s protocol.

#### Establishment of Sec23IP-KO cells

To generate Sec23IP-KOs, pSpCas9(BB)-2A-GFP (pX458) harboring sgRNA#3 (5’-AGGCCCAGGGTTGTAAAGCG-3’) targeting *SEC23IP* was transfected into target cells. The next day, GFP-positive cells were collected by the Duke Cancer Institute Flow Cytometry Core Facility using a BD Diva fluorescence-activated cell sorter, followed by limiting dilution to isolate single-cell-derived clones. Successful Sec23IP knockout was validated by WB.

#### Sample preparation for WB

Cells were washed with phosphate-buffered saline (PBS) twice and collected using cell scrapers, followed by centrifugation at 1,000 × g for 3 min. Cell pellets were lysed with 8 M urea in PBS by sonication, followed by centrifugation at 22,640 × g for 15 min at 4 °C. Supernatant was recovered and the protein concentration was measured using BCA kit (#23225 from ThermoFisher Scientific). Protein concentrations were normalized across samples, mixed with 5 × SDS loading buffer, and incubated at 95 °C for 5 min.

#### IP

For O-GlcNAc analyses, cells were lysed by sonication in buffer containing 20 mM Tris HCl (pH = 7.4), 150 mM NaCl, 1 mM EDTA, 1% Triton X-100, and 0.1% SDS, supplemented with protease inhibitor cocktail (Sigma-Aldrich, P8340, 1:100), 50 µM UDP (Sigma-Aldrich, 94330; OGT inhibitor), and 5 µM PUGNAc (Cayman Chemical, 17151; OGA and lysosomal hexosaminidase inhibitor). Then cell lysates were subjected to centrifugation at 22,640 × g for 15 min at 4 °C. Samples were normalized for protein concentration via BCA, as above. Then, samples were incubated with anti-myc antibody (1 µg per 1 mg total protein) or anti-Sec23IP (from Abcepta; 2 µl per 1 mg total protein) at 4 °C for 1 h, followed by incubation with 30 µl of protein A/G UltraLink resin (ThermoFisher, 53133) at 4 °C overnight. After washing beads three times with lysis buffer, bound proteins were eluted with SDS loading buffer and incubated at 95 °C for 5 min. For detecting protein-protein interactions, cells were lysed by sonication in buffer containing 20 mM Tris HCl (pH = 7.4), 150 mM NaCl, 1 mM EDTA, and 1% Triton X-100, supplemented with protease inhibitor cocktail, 50 µM UDP, and 5 µM PUGNAc. Cell lysates were processed as above.

#### WB

Equal amounts of proteins were loaded in each well and separated by Tris-glycine SDS-PAGE as described (45). For enhanced chemiluminescence (ECL), SDS-PAGE gels were electroblotted onto PVDF membranes (88515, ThermoFisher). The membranes were blocked with blocking buffer [2.5% BSA in Tris-buffered saline containing 0.1% Tween-20 (TBS-T)] for 30 min, followed by incubation with primary antibodies diluted in the same blocking buffer overnight at 4 °C. After washing with TBS-T three times, the membranes were then incubated with secondary antibodies conjugated to HRP, diluted 1:10,000 in blocking buffer for 1 h at room temperature. Membranes were washed with TBS-T three times, and signals were detected via ECL (Genesee Scientific 20-300B) and photographic film (LabScientific, XAR ALF 2025). For quantitative fluorescent IBs, gels were electroblotted onto nitrocellulose membranes (162-115, BioRad). After blocking, incubation with primary antibodies, and washing via the same protocol described above, membranes were incubated with secondary antibodies conjugated to IRDye800, diluted 1:20,000 in blocking buffer for 1 h at room temperature. Membranes were washed with TBS-T three times, followed by washing with TBS to remove residual Tween-20. Signals were detected using a LI-COR Odyssey DLx Imaging System. The following antibodies were used: anti-O-GlcNAc (9D1), 1:1,000; anti-O-GlcNAc (RL2), 1:500; anti-O-GlcNAc (18B10), 1:1,000; anti-myc, 1:1,000; anti-Tubulin, 1:50,000; anti-Sec23IP (sigma), 1:500; anti-Sec31A, 1:1,000; anti-Sec23A, 1:2,000.

#### Collagen transport assay

Kinetics of collagen transport were determined as described previously (47). Briefly, cells were grown on 18-mm coverslips in 24-well plates and pre-incubated at 40 °C in standard DMEM supplemented with 20 mM HEPES-NaOH (pH 7.4) for 3 h. Soon after preincubation, cells for 0 min samples were washed with cold PBS twice and fixed with 4% paraformaldehyde (PFA) at 32 °C for 15 min. Culture medium for the remainder of the samples was replaced with pre-warmed medium supplemented with 20 mM HEPES-NaOH (pH 7.4), 100 µg/ml cycloheximide, and 50 µg/ml freshly dissolved ascorbate, and cells were incubated at 32 °C for 15 min. After transport, cells were washed with cold PBS twice and fixed with 4% PFA at 32 °C for 15 min. After fixation, cells were washed with PBS three times, followed by permeabilization with PBS-T for 10 min at room temperature. After permeabilization, cells were incubated with blocking solution, primary antibodies, and secondary antibodies as described below.

#### E-Cadherin transport assay using RUSH system

RUSH assays to determine the kinetics of E-Cadherin transport were conducted as described (47). Briefly, cells grown on 18-mm coverslips in 24-well plates were transfected with a plasmid encoding the E-Cadherin RUSH probe. The next day, at 70-80% confluences, cells were washed with cold PBS twice and fixed with 4% PFA at 37 °C for 15 min (for 0 min samples). Culture medium for the remainder of the samples was replaced with pre-warmed medium supplemented with 100 µg/ml cycloheximide and 50 µM biotin, and cells were incubated at 37 °C for 20 or 40 min. After transport, cells were washed with cold PBS twice and fixed with 4% PFA at 37 °C for 15 min. After fixation, cells were permeabilized, incubated with blocking solution, primary antibodies, and secondary antibodies as described below.

#### IF imaging

Cells were grown on 18-mm coverslips in 24-well plates. After washing with PBS twice, cells were fixed with cold methanol for 15 min at −20 °C. Cells were washed with PBS three times, followed by incubation with blocking solution (PBS containing 5% goat serum, 0.3% Triton X-100, and 0.05% NaN_3_) for 30 min at room temperature and incubated with primary antibodies diluted in blocking solution at 4 °C overnight. After washing with PBS three times, cells were incubated with secondary antibodies conjugated with Alexa dyes diluted 1:1,000 in blocking solution for 1 h at room temperature. After washing with PBS-T three times, coverslips were mounted on slides with ProLong Diamond anti-fade mounting medium. Images were acquired on a Zeiss 780 fluorescence microscope with a 40×/1.4 NA Oil Plan-Apochromat DIC, (UV) VIS-IR or a 63×/1.4 NA Oil Plan-Apochromat DIC oil immersion objective lens for collagen and E-Cadherin RUSH transport assays or other experiments, respectively. Z-stack images were obtained. Images were processed using FIJI Image J software. The following antibodies were used: mouse anti-myc, 1:1,000; anti-Sec16A, 1:300; anti-GFP, 1:2,000; anti-GM130, 1:500; anti-Sec23IP (sigma), 1:300; rat anti-myc, 1:1,000; anti-Sec31A, 1:300.

#### Quantification of IF data

All images were obtained as z-stacks and processed with Fiji as described previously (47). Before analysis, all images were projected with Max Intensity and subjected to background subtraction with a rolling-ball radius with 50.0 pixels. For collagen transport and E-Cadherin RUSH assays, FIs of GFP were measured to obtain total FIs. Images of Golgi-localized GFP were created by imageCalculator with the “AND create” algorithm with GFP and GM130 images to extract regions of overlap, and FIs of Golgi-localized GFP were measured in them. MFIs in puncta of Sec23IP and Sec31A and the number of Sec23IP and Sec31A puncta were measured in images produced with the appropriate thresholding method with the Analyze Particles plug-in and default settings.

#### Sequence alignment of Sec23IP

Sequence alignment was generated with Clustal Omega and visualized with Boxshade. Amino acids sequences were obtained from Uniprot: *H. sapiens* Sec23IP (Q9Y6Y8-1); *M. musculus* Sec23IP (Q6NZC7); *R. norvegicus* Sec23IP (G3V8Q8); *X. laevis* Sec23IP (A1L2P0); *D. rerio* Sec23IP (Q08BR7).

#### Statistics

Statistical analyses were performed using GraphPad Prism 9 or 10 software (GraphPad Software, Inc.).

## Data availability

All data are contained within the manuscript.

## Supporting information

This article contains supporting information.

## Acknowledgments

We thank Dr. So Young Kim of the Duke Functional Genomics Core for construction of plasmids encoding sgRNAs and Dr. Bin Li of the Duke Cancer Institute Flow Cytometry Shared Resource for cell sorting. We also thank members of the Boyce lab for constructive discussions.

## Supplemental Figure legends

**Figure S1.**
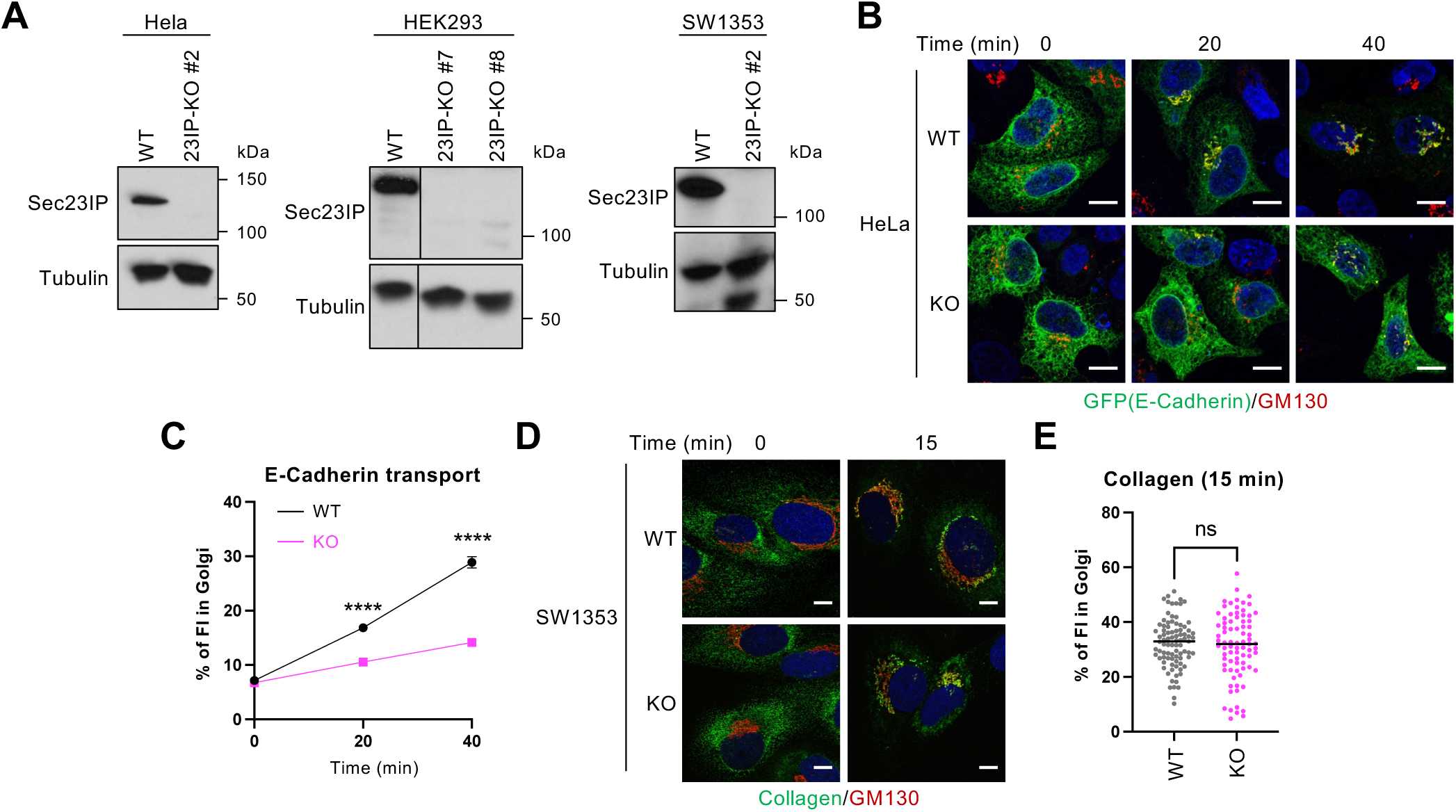
Generation of Sec23IP-KO cells. A. WT and Sec23IP-KO HeLa, HEK293, and SW1353 cells were analyzed by WB. n ≥ 2, representative images shown. Vertical lines in HEK293 blots indicate where irrelevant intervening lanes were removed from the image. B. WT and Sec23IP-KO HeLa cells expressing the E-cadherin RUSH system were incubated with 50 µM biotin and 100 µg/ml cycloheximide for the indicated times and analyzed by IF for E-cadherin (GFP) and GM130. n = 3, representative images shown. Scale bars: 10 µm. C. Quantification of experiments depicted in B. Percentage of GFP FI co-localizing with the Golgi (GM130) was calculated (n = 3). Data were analyzed by Multiple unpaired *t*-test. **** *p* < 0.0001. D. WT and Sec23IP-KO SW1353 cells were incubated at 40 °C for 3 h, shifted to 32 °C plus 50 µg/ml sodium ascorbate and 100 µg/ml cycloheximide for the indicated times, and analyzed by IF for collagen I (collagen) and GM130 (Golgi marker). n = 3, representative images shown. Scale bars: 10 µm. E. Quantification of experiments depicted in D. Percentage of collagen FI co-localizing with the Golgi (GM130) was calculated (n = 3). Data were analyzed by Welch’s *t*-test. ns: not significant.

**Figure S2.**
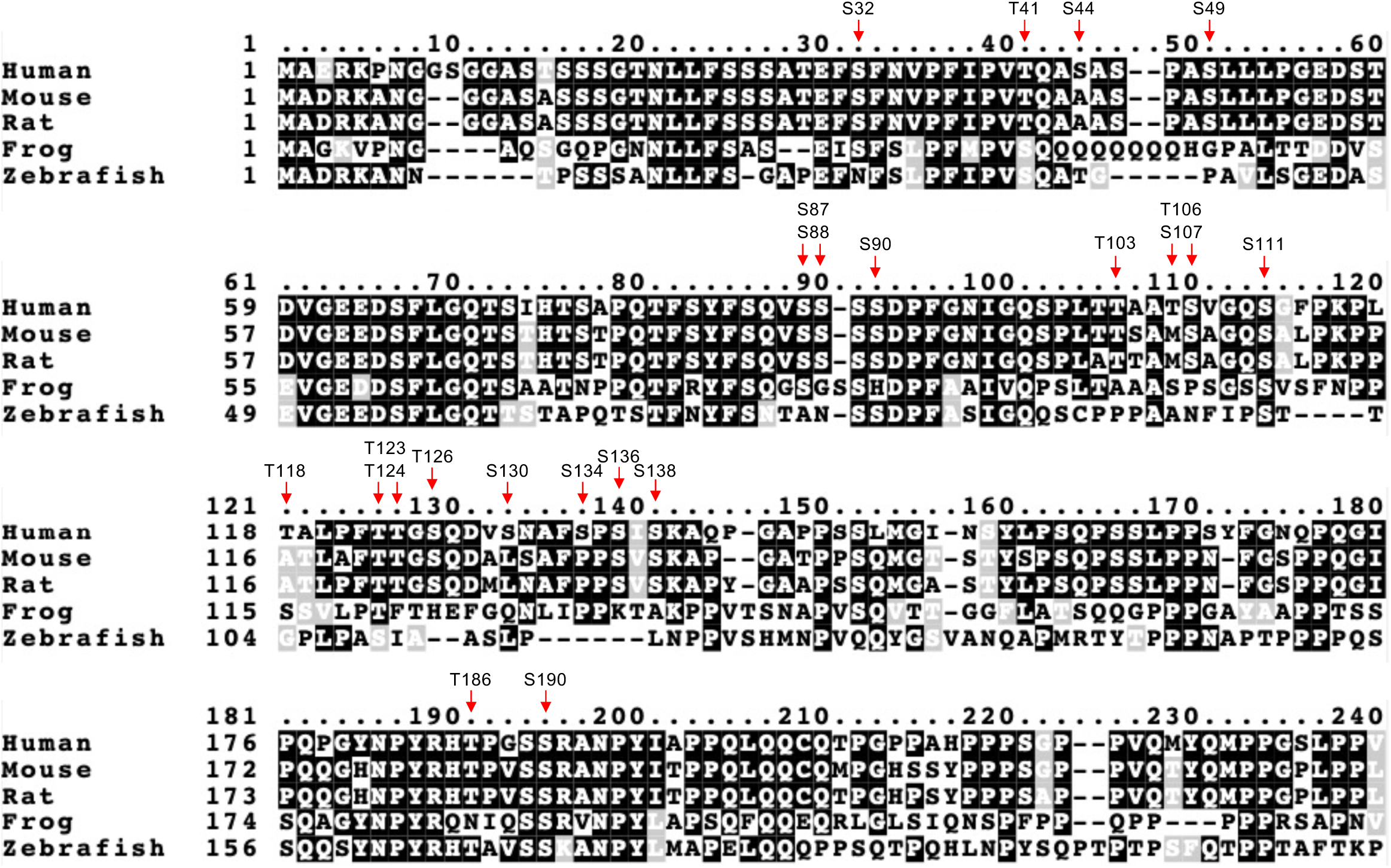
Alignment of representative vertebrate Sec23IP protein sequences. Amino acid sequences of the N-terminal regions of Sec23IP orthologs from human, mouse, rat, frog, and zebrafish were aligned. The human Sec23IP O-GlcNAc sites reported in prior MS studies are indicated by red arrows (see text for references).

**Figure S3.**
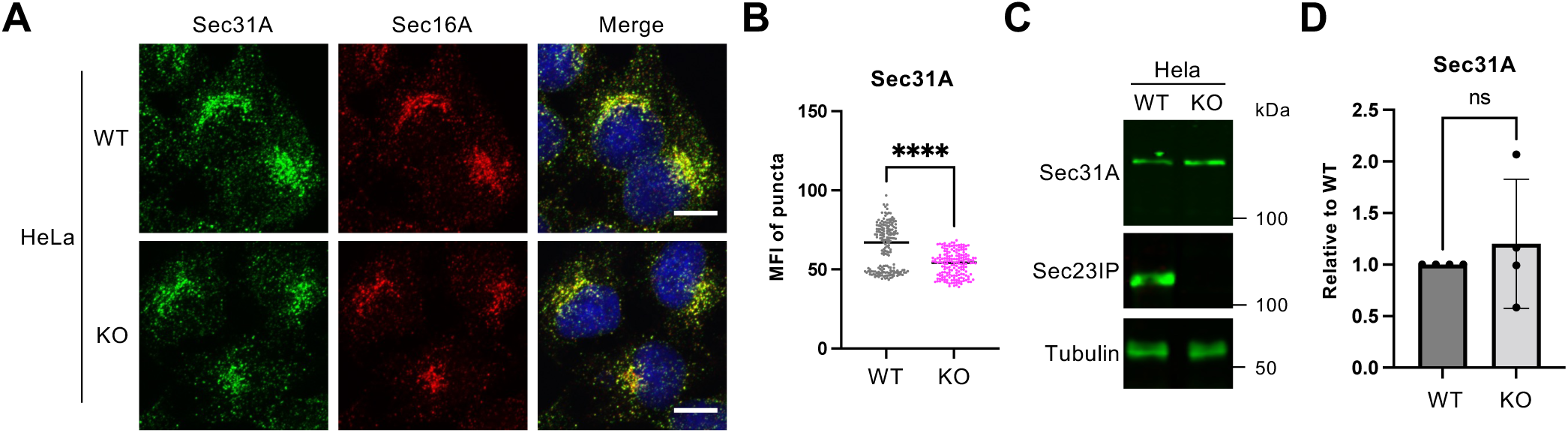
Sec23IP is required for Sec31A recruitment to ERES. A. HeLa-Sec23IP-KO cells were analyzed by IF. n = 3, representative images shown. Sec16A: ERES marker. Scale bars: 10 µm. B. Quantification of experiments depicted in A. MFI of Sec31A puncta was calculated (n = 3). Data were analyzed by Welch’s *t*-test. **** *p* < 0.0001. C. WT and Sec23IP-KO HeLa cells were analyzed by WB. n = 4, representative images shown. D. Quantification of experiments depicted in C. Ratio of Sec31A to tubulin FIs was normalized to WT samples (n = 4). Data were analyzed by Welch’s *t*-test. ns: not significant.

## References

1. Gurkan, C., Stagg, S. M., Lapointe, P., and Balch, W. E. (2006) The COPII cage: unifying principles of vesicle coat assembly Nat Rev Mol Cell Biol 7, 727–738 10.1038/nrm2025

2. Flood, J. R., Mendina, C. A., and Audhya, A. (2025) Organizing principles underlying COPII-mediated transport Curr Opin Cell Biol 94, 102492 10.1016/j.ceb.2025.102492

3. Tang, V. T., and Ginsburg, D. (2023) Cargo selection in endoplasmic reticulum-to-Golgi transport and relevant diseases J Clin Invest 133, 10.1172/JCI163838

4. Robinson, C. M., Duggan, A., and Forrester, A. (2024) ER exit in physiology and disease Front Mol Biosci 11, 1352970 10.3389/fmolb.2024.1352970

5. Fromme, J. C., Ravazzola, M., Hamamoto, S., Al-Balwi, M., Eyaid, W., Boyadjiev, S. A. et al. (2007) The genetic basis of a craniofacial disease provides insight into COPII coat assembly Dev Cell 13, 623–634 10.1016/j.devcel.2007.10.005

6. Garbes, L., Kim, K., Riess, A., Hoyer-Kuhn, H., Beleggia, F., Bevot, A. et al. (2015) Mutations in SEC24D, encoding a component of the COPII machinery, cause a syndromic form of osteogenesis imperfecta Am J Hum Genet 96, 432–439 10.1016/j.ajhg.2015.01.002

7. Liu, Z., Yan, M., Lei, W., Jiang, R., Dai, W., Chen, J., et al. (2022) Sec13 promotes oligodendrocyte differentiation and myelin repair through autocrine pleiotrophin signaling J Clin Invest 132, 10.1172/JCI155096

8. Halperin, D., Kadir, R., Perez, Y., Drabkin, M., Yogev, Y., Wormser, O. et al. (2019) SEC31A mutation affects ER homeostasis, causing a neurological syndrome J Med Genet 56, 139–148 10.1136/jmedgenet-2018-105503

9. Jones, B., Jones, E. L., Bonney, S. A., Patel, H. N., Mensenkamp, A. R., Eichenbaum-Voline, S. et al. (2003) Mutations in a Sar1 GTPase of COPII vesicles are associated with lipid absorption disorders Nat Genet 34, 29–31 10.1038/ng1145

10. Downes, K. W., and Zanetti, G. (2025) Mechanisms of COPII coat assembly and cargo recognition in the secretory pathway Nat Rev Mol Cell Biol 10.1038/s41580-025-00839-y

11. Saxena, S., Foresti, O., Liu, A., Androulaki, S., Pena Rodriguez, M., Raote, I. et al. (2024) Endoplasmic reticulum exit sites are segregated for secretion based on cargo size Dev Cell 59, 2593–2608 e2596 10.1016/j.devcel.2024.06.009

12. Shomron, O., Nevo-Yassaf, I., Aviad, T., Yaffe, Y., Zahavi, E. E., Dukhovny, A., et al. (2021) COPII collar defines the boundary between ER and ER exit site and does not coat cargo containers J Cell Biol 220, 10.1083/jcb.201907224

13. Weigel, A. V., Chang, C. L., Shtengel, G., Xu, C. S., Hoffman, D. P., Freeman, M. et al. (2021) ER-to-Golgi protein delivery through an interwoven, tubular network extending from ER Cell 184, 2412–2429 e2416 10.1016/j.cell.2021.03.035

14. Downes, K., Flood, J., Nans, A., Van der Verren, S., Audhya, A., and Zanetti, G. (2025) Multi-scale Molecular Imaging of Human Cells reveals COPI and COPII Vesicles at ER Exit Sites bioRxiv 2025.2007.2029.667472 10.1101/2025.07.29.667472

15. Matsuoka, K., Orci, L., Amherdt, M., Bednarek, S. Y., Hamamoto, S., Schekman, R. et al. (1998) COPII-coated vesicle formation reconstituted with purified coat proteins and chemically defined liposomes Cell 93, 263–275 10.1016/s0092-8674(00)81577-9

16. Zanetti, G., Prinz, S., Daum, S., Meister, A., Schekman, R., Bacia, K., et al. (2013) The structure of the COPII transport-vesicle coat assembled on membranes Elife 2, e00951 10.7554/eLife.00951

17. Khoriaty, R., Hesketh, G. G., Bernard, A., Weyand, A. C., Mellacheruvu, D., Zhu, G. et al. (2018) Functions of the COPII gene paralogs SEC23A and SEC23B are interchangeable in vivo Proc Natl Acad Sci U S A 115, E7748–E7757 10.1073/pnas.1805784115

18. Tang, V. T., Xiang, J., Chen, Z., McCormick, J., Abbineni, P. S., Chen, X. W., et al. (2024) Functional overlap between the mammalian Sar1a and Sar1b paralogs in vivo Proc Natl Acad Sci U S A 121, e2322164121 10.1073/pnas.2322164121

19. Watson, P., Townley, A. K., Koka, P., Palmer, K. J., and Stephens, D. J. (2006) Sec16 defines endoplasmic reticulum exit sites and is required for secretory cargo export in mammalian cells Traffic 7, 1678–1687 10.1111/j.1600-0854.2006.00493.x

20. Bi, X., Corpina, R. A., and Goldberg, J. (2002) Structure of the Sec23/24-Sar1 pre-budding complex of the COPII vesicle coat Nature 419, 271–277 10.1038/nature01040

21. Gimeno, R. E., Espenshade, P., and Kaiser, C. A. (1996) COPII coat subunit interactions: Sec24p and Sec23p bind to adjacent regions of Sec16p Mol Biol Cell 7, 1815–1823 10.1091/mbc.7.11.1815

22. Townley, A. K., Feng, Y., Schmidt, K., Carter, D. A., Porter, R., Verkade, P. et al. (2008) Efficient coupling of Sec23-Sec24 to Sec13-Sec31 drives COPII-dependent collagen secretion and is essential for normal craniofacial development J Cell Sci 121, 3025–3034 10.1242/jcs.031070

23. Wendeler, M. W., Paccaud, J. P., and Hauri, H. P. (2007) Role of Sec24 isoforms in selective export of membrane proteins from the endoplasmic reticulum EMBO Rep 8, 258–264 10.1038/sj.embor.7400893

24. Ong, Y. S., Tang, B. L., Loo, L. S., and Hong, W. (2010) p125A exists as part of the mammalian Sec13/Sec31 COPII subcomplex to facilitate ER-Golgi transport J Cell Biol 190, 331–345 10.1083/jcb.201003005

25. Tani, K., Mizoguchi, T., Iwamatsu, A., Hatsuzawa, K., and Tagaya, M. (1999) p125 is a novel mammalian Sec23p-interacting protein with structural similarity to phospholipid-modifying proteins J Biol Chem 274, 20505–20512 10.1074/jbc.274.29.20505

26. Nakajima, K., Sonoda, H., Mizoguchi, T., Aoki, J., Arai, H., Nagahama, M., et al. (2002) A novel phospholipase A1 with sequence homology to a mammalian Sec23p-interacting protein, p125 J Biol Chem 277, 11329–11335 10.1074/jbc.M111092200

27. Klinkenberg, D., Long, K. R., Shome, K., Watkins, S. C., and Aridor, M. (2014) A cascade of ER exit site assembly that is regulated by p125A and lipid signals J Cell Sci 127, 1765–1778 10.1242/jcs.138784

28. Du, Y., Fan, X., Song, C., Chang, W., Xiong, J., Deng, L. et al. (2024) Sec23IP recruits VPS13B/COH1 to ER exit site-Golgi interface for tubular ERGIC formation J Cell Biol 223, 10.1083/jcb.202402083

29. Long, K. R., Singh, G., Villemeur, M., Brouwers, N., Malhotra, V., Raote, I., et al. (2025) p125A (Sec23ip) couples COPII coat assembly with donor-acceptor membrane organization to facilitate tunnel-based traffic bioRxiv 10.1101/2025.05.07.652703

30. Woo, C. M., Lund, P. J., Huang, A. C., Davis, M. M., Bertozzi, C. R., and Pitteri, S. J. (2018) Mapping and Quantification of Over 2000 O-linked Glycopeptides in Activated Human T Cells with Isotope-Targeted Glycoproteomics (Isotag) Mol Cell Proteomics 17, 764–775 10.1074/mcp.RA117.000261

31. Liu, J., Hao, Y., Wang, C., Jin, Y., Yang, Y., Gu, J. et al. (2022) An Optimized Isotopic Photocleavable Tagging Strategy for Site-Specific and Quantitative Profiling of Protein O-GlcNAcylation in Colorectal Cancer Metastasis ACS Chem Biol 17, 513–520 10.1021/acschembio.1c00981

32. Kim, D. I., Cutler, J. A., Na, C. H., Reckel, S., Renuse, S., Madugundu, A. K. et al. (2018) BioSITe: A Method for Direct Detection and Quantitation of Site-Specific Biotinylation J Proteome Res 17, 759–769 10.1021/acs.jproteome.7b00775

33. Qin, K., Zhu, Y., Qin, W., Gao, J., Shao, X., Wang, Y. L. et al. (2018) Quantitative Profiling of Protein O-GlcNAcylation Sites by an Isotope-Tagged Cleavable Linker ACS Chem Biol 13, 1983–1989 10.1021/acschembio.8b00414

34. Wang, S., Yang, F., Petyuk, V. A., Shukla, A. K., Monroe, M. E., Gritsenko, M. A. et al. (2017) Quantitative proteomics identifies altered O-GlcNAcylation of structural, synaptic and memory-associated proteins in Alzheimer’s disease J Pathol 243, 78–88 10.1002/path.4929

35. Bond, M. R., and Hanover, J. A. (2015) A little sugar goes a long way: the cell biology of O-GlcNAc J Cell Biol 208, 869–880 10.1083/jcb.201501101

36. Yang, X., and Qian, K. (2017) Protein O-GlcNAcylation: emerging mechanisms and functions Nat Rev Mol Cell Biol 18, 452–465 10.1038/nrm.2017.22

37. King, D. T., Males, A., Davies, G. J., and Vocadlo, D. J. (2019) Molecular mechanisms regulating O-linked N-acetylglucosamine (O-GlcNAc)-processing enzymes Curr Opin Chem Biol 53, 131–144 10.1016/j.cbpa.2019.09.001

38. Wulff-Fuentes, E., Berendt, R. R., Massman, L., Danner, L., Malard, F., Vora, J. et al. (2021) The human O-GlcNAcome database and meta-analysis Sci Data 8, 25 10.1038/s41597-021-00810-4

39. Hart, G. W. (2014) Three Decades of Research on O-GlcNAcylation - A Major Nutrient Sensor That Regulates Signaling, Transcription and Cellular Metabolism Front Endocrinol (Lausanne) 5, 183 10.3389/fendo.2014.00183

40. Bisnett, B. J., Condon, B. M., Lamb, C. H., Georgiou, G. R., and Boyce, M. (2020) Export Control: Post-transcriptional Regulation of the COPII Trafficking Pathway Front Cell Dev Biol 8, 618652 10.3389/fcell.2020.618652

41. Ye, L., Ding, W., Xiao, D., Jia, Y., Zhao, Z., Ao, X. et al. (2023) O-GlcNAcylation: cellular physiology and therapeutic target for human diseases MedComm (2020) 4, e456 10.1002/mco2.456

42. Fahie, K. M. M., Papanicolaou, K. N., and Zachara, N. E. (2022) Integration of O-GlcNAc into Stress Response Pathways Cells 11, 10.3390/cells11213509

43. Le Minh, G., Esquea, E. M., Young, R. G., Huang, J., and Reginato, M. J. (2023) On a sugar high: Role of O-GlcNAcylation in cancer J Biol Chem 299, 105344 10.1016/j.jbc.2023.105344

44. Bisnett, B. J., Condon, B. M., Linhart, N. A., Lamb, C. H., Huynh, D. T., Bai, J. et al. (2021) Evidence for nutrient-dependent regulation of the COPII coat by O-GlcNAcylation Glycobiology 31, 1102–1120 10.1093/glycob/cwab055

45. Cox, N. J., Unlu, G., Bisnett, B. J., Meister, T. R., Condon, B. M., Luo, P. M. et al. (2018) Dynamic Glycosylation Governs the Vertebrate COPII Protein Trafficking Pathway Biochemistry 57, 91–107 10.1021/acs.biochem.7b00870

46. Cho, H. J., and Mook-Jung, I. (2018) O-GlcNAcylation regulates endoplasmic reticulum exit sites through Sec31A modification in conventional secretory pathway FASEB J 32, 4641–4657 10.1096/fj.201701523R

47. Hirata, T., Choudhary, D., Bisnett, B. J., Soderblom, E. J., Knapik, E. W., and Boyce, M. (2025) Dynamic regulation of the COPII interactome and collagen trafficking by site-specific glycosylation of Sec24D bioRxiv 2025.2006.2013.659590 10.1101/2025.06.13.659590

48. Georgiou, G. R., Hirata, T., Soderblom, E., Homoelle, R., Maiwald, J., and Boyce, M. (2025) Dynamic regulation of Sec24C by phosphorylation and O-GlcNAcylation during cell cycle progression J Biol Chem 110456 10.1016/j.jbc.2025.110456

49. Gloster, T. M., Zandberg, W. F., Heinonen, J. E., Shen, D. L., Deng, L., and Vocadlo, D. J. (2011) Hijacking a biosynthetic pathway yields a glycosyltransferase inhibitor within cells Nat Chem Biol 7, 174–181 10.1038/nchembio.520

50. Yuzwa, S. A., Macauley, M. S., Heinonen, J. E., Shan, X., Dennis, R. J., He, Y. et al. (2008) A potent mechanism-inspired O-GlcNAcase inhibitor that blocks phosphorylation of tau in vivo Nat Chem Biol 4, 483–490 10.1038/nchembio.96

51. Teo, C. F., Ingale, S., Wolfert, M. A., Elsayed, G. A., Not, L. G., Chatham, J. C. et al. (2010) Glycopeptide-specific monoclonal antibodies suggest new roles for O-GlcNAc Nat Chem Biol 6, 338–343 10.1038/nchembio.338

52. Nishimura, N., and Balch, W. E. (1997) A di-acidic signal required for selective export from the endoplasmic reticulum Science 277, 556–558 10.1126/science.277.5325.556

53. Subramanian, A., Capalbo, A., Iyengar, N. R., Rizzo, R., di Campli, A., Di Martino, R. et al. (2019) Auto-regulation of Secretory Flux by Sensing and Responding to the Folded Cargo Protein Load in the Endoplasmic Reticulum Cell 176, 1461–1476 e1423 10.1016/j.cell.2019.01.035

54. Boncompain, G., Divoux, S., Gareil, N., de Forges, H., Lescure, A., Latreche, L. et al. (2012) Synchronization of secretory protein traffic in populations of cells Nat Methods 9, 493–498 10.1038/nmeth.1928

55. Vincourt, J. B., Etienne, S., Cottet, J., Delaunay, C., Malanda, C. B., Lionneton, F. et al. (2010) C-propeptides of procollagens I alpha 1 and II that differentially accumulate in enchondromas versus chondrosarcomas regulate tumor cell survival and migration Cancer Res 70, 4739–4748 10.1158/0008-5472.CAN-10-0046

56. Mironov, A. A., Beznoussenko, G. V., Nicoziani, P., Martella, O., Trucco, A., Kweon, H. S. et al. (2001) Small cargo proteins and large aggregates can traverse the Golgi by a common mechanism without leaving the lumen of cisternae J Cell Biol 155, 1225–1238 10.1083/jcb.200108073

57. Ben Ahmed, A., Lemaire, Q., Scache, J., Mariller, C., Lefebvre, T., and Vercoutter-Edouart, A. S. (2023) O-GlcNAc Dynamics: The Sweet Side of Protein Trafficking Regulation in Mammalian Cells Cells 12, 10.3390/cells12101396

58. Banani, S. F., Lee, H. O., Hyman, A. A., and Rosen, M. K. (2017) Biomolecular condensates: organizers of cellular biochemistry Nat Rev Mol Cell Biol 18, 285–298 10.1038/nrm.2017.7

59. Molliex, A., Temirov, J., Lee, J., Coughlin, M., Kanagaraj, A. P., Kim, H. J. et al. (2015) Phase separation by low complexity domains promotes stress granule assembly and drives pathological fibrillization Cell 163, 123–133 10.1016/j.cell.2015.09.015

60. Elbaum-Garfinkle, S., Kim, Y., Szczepaniak, K., Chen, C. C., Eckmann, C. R., Myong, S. et al. (2015) The disordered P granule protein LAF-1 drives phase separation into droplets with tunable viscosity and dynamics Proc Natl Acad Sci U S A 112, 7189–7194 10.1073/pnas.1504822112

61. Burke, K. A., Janke, A. M., Rhine, C. L., and Fawzi, N. L. (2015) Residue-by-Residue View of In Vitro FUS Granules that Bind the C-Terminal Domain of RNA Polymerase II Mol Cell 60, 231–241 10.1016/j.molcel.2015.09.006

62. Wang, X., Huang, R., Wang, Y., Zhou, W., Hu, Y., Yao, Y. et al. (2023) Manganese regulation of COPII condensation controls circulating lipid homeostasis Nat Cell Biol 25, 1650–1663 10.1038/s41556-023-01260-3

63. Gallo, R., Rai, A. K., McIntyre, A. B. R., Meyer, K., and Pelkmans, L. (2023) DYRK3 enables secretory trafficking by maintaining the liquid-like state of ER exit sites Dev Cell 58, 1880–1897 e1811 10.1016/j.devcel.2023.08.005

64. Johnson, A., Bhattacharya, N., Hanna, M., Pennington, J. G., Schuh, A. L., Wang, L. et al. (2015) TFG clusters COPII-coated transport carriers and promotes early secretory pathway organization EMBO J 34, 811–827 10.15252/embj.201489032

65. Lv, P., Du, Y., He, C., Peng, L., Zhou, X., Wan, Y. et al. (2022) O-GlcNAcylation modulates liquid-liquid phase separation of SynGAP/PSD-95 Nat Chem 14, 831–840 10.1038/s41557-022-00946-9

66. Nosella, M. L., Tereshchenko, M., Pritisanac, I., Chong, P. A., Toretsky, J. A., Lee, H. O., et al. (2021) O-Linked-N-Acetylglucosaminylation of the RNA-Binding Protein EWS N-Terminal Low Complexity Region Reduces Phase Separation and Enhances Condensate Dynamics J Am Chem Soc 143, 11520–11534 10.1021/jacs.1c04194

67. Li, X., Pinou, L., Du, Y., Chen, X., and Liu, C. (2023) Emerging roles of O-glycosylation in regulating protein aggregation, phase separation, and functions Curr Opin Chem Biol 75, 102314 10.1016/j.cbpa.2023.102314

68. Xu, S., Yin, K., Xu, X., Fu, L., and Wu, R. (2025) O-GlcNAcylation reduces proteome solubility and regulates the formation of biomolecular condensates in human cells Nat Commun 16, 4068 10.1038/s41467-025-59371-4

